# Cross-Linking of T cell to B cell lymphoma by the T cell bispecific antibody CD20-TCB induces IFNγ/CXCL10-dependent peripheral T cell recruitment in humanized murine model

**DOI:** 10.1101/2020.10.09.332874

**Authors:** Floriana Cremasco, Elena Menietti, Dario Speziale, Johannes Sam, Stefano Sammicheli, Marine Richard, Ahmet Varol, Christian Klein, Pablo Umana, Marina Bacac, Sara Colombetti, Mario Perro

## Abstract

Diffuse large B cell lymphomas (DLBCL) are a highly heterogeneous subtype of Non Hodgkin Lymphoma (NHL), accounting for about 25% of NHL [1]. Despite an increased progression-free survival upon therapy, 40-50% of patients develop relapse/refractory disease, therefore there remains an important medical need [2]. T cell recruiting therapies, such as the CD20xCD3 T cell bi-specific antibody CD20-TCB (RG6026 or glofitamab), represent a novel approach to target all stages of DLBCL, especially those that fail to respond to multiple lines of treatment [3, 4]. We aimed for a better understanding of the molecular features related to the mode of action (MoA) of CD20-TCB in inducing Target/T cell synapse formation and human T cell recruitment to the tumor. To directly evaluate the correlation between synapse, cytokine production and anti-tumor efficacy using CD20-TCB, we developed an innovative preclinical human DLBCL *in vivo* model that allowed tracking *in vivo* human T cell dynamics by multiphoton intravital microscopy (MP-IVM). By *ex vivo* and *in vivo* approaches, we revealed that CD20-TCB is inducing strong and stable synapses between human T cell and tumor cells, which are dependent on the dose of CD20-TCB and on LFA-1 activity but not on FAS-L. Moreover, despite CD20-TCB being a large molecule (194.342 kDa), we observed that intra-tumor CD20-TCB*-*mediated human T cell-tumor cell synapses occur within 1 hour upon CD20-TCB administration. These tight interactions, observed for at least 72 hours post TCB administration, result in tumor cell cytotoxicity, resident T cell proliferation and peripheral blood T cell recruitment into tumor. By blocking the IFNγ-CXCL10 axis, the recruitment of peripheral T cells was abrogated, partially affecting the efficacy of CD20-TCB treatment which rely only on resident T cell proliferation. Altogether these data reveal that CD20-TCB’s anti-tumor activity relies on a triple effect: i) fast formation of stable T cell-tumor cell synapses which induce tumor cytotoxicity and cytokine production, ii) resident T cell proliferation and iii) recruitment of fresh peripheral T cells to the tumor core to allow a positive enhancement of the anti-tumor effect.

## Introduction

Diffuse large B-cell lymphoma (DLBCL) is the most common histological subtype of B-cell NHL accounting for 25-30% of diagnosed NHL [5, 6]. The majority of DLBCL lymphomas occur in the lymph nodes, however about 30% of them occur in extranodal sites. Although heterogeneous, all DLBCL, including the extranodal types, share the characteristic expression of CD19 and CD20 on tumor cells. As a consequence, since the discovery of rituximab, a chimeric monoclonal antibody against CD20, the clinical practice standard of care for all forms of DLBCL is the combination of rituximab with chemotherapy (R-CHOP) [7, 8]. However, patients who fail to respond to first line treatment, or that develop resistance (40-50%), have poor prognosis and DLBCL still represents a major medical need.

Currently, the use of T cell bi-specific antibodies targeting CD20 on tumor cells, e.g. CD20-TCB (also known as RG6026 or glofitamab), is a promising approach for the treatment of DLBCL [3, 4]. Indeed, CD20-TCB can redirect the activity of conventional CD4 and CD8 T cells against lymphoma cells by concomitant binding of CD20 on tumor cells and CD3 on T cells [3, 9]. Previous studies revealed that CD20-TCB has a long half-life and induces selective activation of conventional T cells, resulting in efficient tumor regression of the aggressive DLBCL xenograft model WSU DLCL2 in preclinical animal models in human stem cell humanized (HSC) NSG mice [3].Tumor regression has been also observed in several clinical trials of patient with r/r DLBCL [10-13]. The treatment with such a potent CD20-TCB increases the risk of cytokine release syndrome (CRS) due to the peripheral cross-linking between T cell and B cells. Therefore, CD20-TCB administration is preceded by a single administration of Obinutuzumab (Gazyva), as a pretreatment (Gpt) to deplete circulating CD20^+^ cells and thus improve the safety of the first CD20-TCB administration [3]. Altogether, Gpt plus CD20-TCB represents a potent approach for the treatment of lymphoma patients and is currently being evaluated in multicenter Phase I/Ib studies in individuals with relapsed/refractory NHL (NCT03075696, NCT03467373, NCT03533283, NCT04077723). It has been demonstrated that clinical efficacy of checkpoint inhibitors (CPIs) positively correlates with the degree of conventional CD4 and CD8 T cell infiltration in the so called “inflamed” tumors, while tumors that are devoid of T cell infiltration (“immune desert”) or that are only partially infiltrated (“immune excluded”), are generally poor responders to immunotherapies [14-16]. Interestingly, TCBs have shown to be efficacious even in tumors with poor immune cell infiltration and to convert, upon treatment, immune excluded and desert tumors into inflamed ones [3, 17, 18]. In view of the promising efficacy, it’s important to elucidate the molecular features of CD20-TCB activity, including: 1) how quickly CD20-TCB binds to tumor cells and induces T cell-target cells synapse formation; 2) which are the features of CD20-TCB mediated synapses; 3) for how long can CD20-TCB mediate T cell engagement to tumor cells; 4) which are the molecular drivers that turn a desert tumor into an inflamed one after CD20-TCB treatment.

We took advantage of both *in vitro* and *in vivo* approaches. Live confocal *in vitro* imaging allowed in-depth description of CD20-TCB-mediated T cell-tumor cell synapses. In order to visualize the intra-tumor response of the human immune system to CD20-TCB *in vivo*, we imaged using multiphoton intravital microscopy (MP-IVM) a newly set-up model of humanized mice with reduced xenoreaction that allow better analysis of T cell in vivo dynamics compare to previous models [19-21]. Using this technique, we were able to assess the kinetics and MoA of CD20-TCB *in vivo*, as well as highlighting the kinetics of TCB-induced cytotoxicity and identifying the mechanisms involved in CD20-TCB-dependent peripheral human T cell recruitment to tumors.

## Materials and Methods

### Cell culture and labelling

WSU DLCL2 cells (human diffuse large B cell lymphoma) were originally obtained from ECACC (European Collection of Cell Culture) and deposited after expansion in the Roche Glycart internal cell bank. Cells were cultured in DMEM containing 10% FCS.

OCI-Ly18 cells (human diffuse large B cell lymphoma) were originally obtained from Deutsche Sammlung von Mikroorganismen und Zellkulturen GmbH (DSMZ) and deposited after expansion in the Roche Glycart internal cell bank. Cells were cultivated in RPMI 1640 medium containing 10% FCS and 1% Glutamax. CT26 cells (*Mus Musculus* colon carcinoma) were originally obtained from ATCC and after expansion deposited in the Roche Glycart internal cell bank, and were cultured in RPMI containing 10% FCS and 1% Glutamax.

All cells were cultured at 37 °C in a water-saturated atmosphere at 5 % CO_2_. All cell lines were tested negative to mycoplasma by PCR assay. Conventional pan T cells from the spleen of HSC-NSG mice were isolated using a CD2^+^ selection kit (Miltenyi Biotech, 130-091-114) and directly injected.

Conventional pan T cells or CD8^+^ T cells from peripheral blood mononuclear cells (PBMCs) were isolated from buffy coats (Blutspende Zurich) by negative magnetic isolation using Pan T cell isolation kit II (Miltenyi Biotech, 130-096-535) or CD8^+^ T cell isolation kit II (Miltenyi Biotech, 130-096-495), respectively. For *in vitro* activation, CD8^+^ T cells were cultivated at 1.5×10^6^ cells/ml in CST™ OpTimizer T cell expansion SFM medium (Thermo Fisher, A1048501) supplemented with 20 U/ml of recombinant huIL-2 (Roche, 11147528001) and then plated into 12 well plates previously coated for 2 hours RT with 1 μg/ml of huCD28.2 (BioLegend) and 1 μg/ml of human CD3ε (BioLegend). When needed, cells were fluorescently labeled with blue (10 μM CellTracker™ Blue CMAC Dye,, Thermo Fisher Scientific, C2110), red (4 μM CellTracker™ Orange CMTMR Dye, Thermo Fisher Scientific, C2927) or green (2.5 μM CellTracker™ Green CMFDA Dye, Thermo Fisher Scientific, C2925), immediately before injection. Labeling was performed at 37 °C for 20 minutes in RPMI medium without serum. When required, labeling with 1.25 μM of CellTrace™CFSE Cell proliferation kit (Thermo Fisher Scientific, C34554) has been performed in PBS at 37 °C for 20 minutes.

### Flow cytometry

Blood from HSC-NSG mice, PBMC-NSG mice or from human PBMCs was collected and lysed using BD Pharm Lyse (Cat. 555899). The blood lysates were then stained with the following panels: 1) muCD45 AF488 (BioLegend), huCD45 APCCy7 (BioLegend), huCD3 BV605 (BioLegend), huCD19 AF647 (BioLegend), huCD33 PerCp-Cy5.5 (BioLegend), NKp46 BV421 (BioLegend); 2) huCD45 APC Cy7 (BioLegend), huCD3 PerCp-Cy5.5 (BioLegend), GZB PE (eBiosciences), Ki-67 AF647 (BD); 3) CD62L APC-Cy7 (BioLegend), huCD45 PE (BioLegend), huCD3 BV605 (BioLegend), CD45RA BV711 (BioLegend).

For the evaluation of killing capacity, cells were stained with huCD8 AF700 (BioLegend) followed by Annexin V-PI staining according to manufacturer’s instruction (BioLegend, APC Annexin V apoptosis detection kit with PI, 640932).

Characterization of T cell activation has been evaluated with extracellular markers including: huCD3 BV605 (BioLegend), huCD8 BV510 (BioLegend), huCD8 AF700 (BioLegend), huCD4 AF488 (BioLegend), huCD69 BV421 (BioLegend), huCD69 AF647 (BioLegend), huCD25 BV711 (BioLegend). In order to discriminate between Live and Dead cells we always included Fixable Viability Dye eFlour™ 780 (eBioscience™). When required cells were incubated for 4 hours with GolgiStop (BD) and GolgiPlug (BD), in order to allow intracellular cytokines accumulation, then cells were fixed and permeabilized according to the manufacturer’s instruction (Fixation and Permeabilization Solution BD, 554714). Staining of intracellular cytokines has been performed with: huIFNγ BV510 (BioLegend), huTNFα PE-Cy7 (BioLegend), huGZB FITC (BioLegend). Cells were run on a BD 5 laser LSR Fortessa. Data were analyzed using FlowJo V.10.

### *In vitro* confocal Imaging

For Live Confocal Imaging 200.000 WSU DLCL2 or OCI-Ly18 cells were stained with 10 μM of CellTracker™ Blue CMAC Dye and then co-plated with 150.000 3T3 NIH Fibroblasts in 8-well plastic-treated bottom slides (Ibidi, 80826). Cells were then incubated overnight at 37 °C. Before acquisition, CD3/CD28-activated CD8^+^ T cells were stained with 2.5 μM of CellTracker™ Green CMFDA Dye and 40.000 labeled T cells were added to tumor cells. Alexa Fluor647-labeled CD20-TCB was added directly into the imaging wells at the indicated concentrations. Slides were imaged with a confocal microscope (inverted LSM 700, Zeiss) with a temperature and CO_2_-controlled stage. Live acquisition was performed with a 20x objective. Movies were collected using Zen software (Zeiss) coupled to the microscope, analyzed with Imaris (Bitplane; Oxford Instruments), and plotted using GraphPad Prism. For imaging on fixed samples, 500.000 WSU DLCL2 cells, stained with 10 μM of CellTracker™ Blue CMAC Dye were plated on glass disc for imaging previously coated with Retronectin (5 μg/cm^2^). After 3h 150.000 CD3-CD28-activated CD8^+^ T cells, were added to the tumor cells and incubated for 2 hours with CD20-TCB (200 ng/ml) and +/- αLFA1 antibody (Clone: TS 1/18, 10 μg/ml, BioLegend, 302116). Cells were then gently washed, fixed and permeabilized according to manufacturer’s instruction (Fixation and Permeabilization Solution BD, 554714). Non specific binding sites were saturated with 40 minutes incubation in PBS + 1% BSA. Cells were then stained with 0.2 μM of Phalloidin AlexaFluor488 (A12379, Thermo Fisher Scientific) for 30min followed by staining with αLFA1-Alexa Fluor647 (Clone: CBR LFA 1/2, BioLegend, 366311) for 30 minutes. Slides were then mounted with FluorMount G (Southern Biotech, 0100-01) and imaged with a confocal microscope (inverted LSM 700, Zeiss). Acquisition was performed with a 20x objective. Images were then analyzed with Imaris (Bitplane; Oxford Instruments).

### *In vitro* killing assays

100.000 WSU DLCL2 or OCI-Ly18 cells were plated together with CD3/CD28-activated CD8^+^ T cells or with freshly purified CD8^+^ T cells from PBMCs at the indicated ratios (1:1; 4:1; 40:1), into 96-U bottom-wells plates. CD20-TCB has been added at the indicated concentrations. When required, wells were supplemented with 10 μg/ml of αLFA1 antibody (Clone: TS 1/18, BioLegend, 302116) or with 10 μg/ml of αFAS-L antibody (BD, 556372). Cells were then incubated at 37 °C in a water-saturated atmosphere at 5 % CO_2_, and analyzed at the indicated time-points. When needed, supernatants were collected and stored at -20 °C and subsequently used for multiplexing analysis.

### Chemotaxis assays

Supernatants freshly derived from killing assay were immediately adopted for chemotaxis assays. 600 μl of supernatant were placed in the bottom chamber of Corning^®^ Transwell^®^ polycarbonate membrane cell culture inserts 6.5 mm Transwell with 5.0 μm pore polycarbonate membrane insert (CLS3421). CD3-CD28-activated CD8^+^ T cells were labeled with 2.5 μM of CellTracker™ Green CMFDA Dye, then resuspended at 1*10^6^ cells/ml in CST™ OpTimizer T cell expansion SFM medium (Thermo Fisher, A1048501) without serum. 100 μl of cell suspension was then seeded in the upper chamber of the transwell. After 3 hours of migration, cells migrated to the bottom chamber were collected and stained with CD8 AF700 (BioLegend) and resuspended in 200 μl of PBS. 150 μl of cell suspension was then acquired at constant flow rate (2.5 μl/sec) on a BD 5 laser LSR Fortessa. Data were analyzed using FlowJo V.10. % of migrated cells has been calculated by applying this formula: 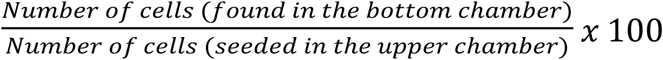. We then calculated the fold change of migration as compared to the untreated conditions (spontaneous migration).

### Multiplex

100.000 WSU DLCL2 cells were seeded in DMEM + 10% FCS + 1% Glutamax, into 96-U bottom-wells plates, and treated with huIFNγ at the indicated concentrations for 24h. Cell culture supernatant was stored at -20 °C and subsequently analyzed together with supernatant derived from killing assays with the Bio-Plex Pro(tm) Human Chemokine Panel, 40-Plex (#171AK99MR2) following manufacturer’s instructions. Samples were then run on Bio-Plex® 200 System. Data were plotted and analyzed with GraphPad Prism.

### *In vivo* studies in mice

All animal protocols were approved by the Cantonal Veterinary Office in Zürich (license ZH193/2014). Mice were NSG (Taconic) mice, aged 10-14 weeks at start of experiments and maintained under specific pathogen free condition with daily cycles of 12 h light and 12 h darkness according to committed guidelines (GV-Solas, Felasa, TierschG). After arrival, animals were maintained for one week to get accustomed to their new environment and for observation. Continuous health monitoring was carried out on a regular basis. For all experiments we adopted 25 g (+/- 5 g) female NSG mice. Mice included in the study were allocated into groups after randomization with a dedicated software.

### Skinfold chamber model

The skinfold chamber surgery was performed according to previously described procedures [22]. NSG mice were anesthetized using isoflurane (1-5 %) in combination with an air-oxygen mixture, and placed on a heating pad. Mice were then shaved with a razor followed by hair-removal cream admixed 50:50 with hydrating cream. After the formation of a dorsal skin fold, a round piece of skin of approximately 10 mm diameter was removed from the first layer in the fold, as well as the superficial layer of fat underneath with the visual aid of a stereomicroscope, exposing the vasculature on the opposite side. The skinfold chamber (APJ trading) was then fixed to the animal with screws pierced through the skin, and secured by suturing the upper edge. Finally, the exposed skin layer was rinsed with saline solution and a coverslip was fixed onto the chamber by a spanning ring. All procedures were performed under a sterile hood in sterile conditions. Mice were allowed to wake under a warming infrared lamp, and provided with a painkiller for 48h. After at least 48h recovery time, mice were anesthetized again by isoflurane, and the coverslip removed. With the visual aid of a stereomicroscope, the desired cell mixture for each experiment was then injected intradermally, 2 injections of 25 μl per mouse, together with the treatment stated in the figure legends, and the coverslip replaced. Mice were then allowed to recover under an infrared lamp, and eventual intravenous administration of drugs was performed concomitantly.

### Intravital 2 photon Imaging

2 photon microscopy was performed on a Zeiss 710 combined confocal and multiphoton system using a 20x objective (Plan Apochromat, Zeiss, N/A=1). Mice were anesthetized by isoflurane (1-5 %) in combination with an air-oxygen mixture, and kept at a constant temperature of 37 °C by using an *ad-hoc* built incubator. By adopting an excitation wavelength of 770nm, with a laser power of 16, we achieved simultaneous excitation of the dyes. Sequential 3D stacks of 40 μm at 4 μm step sizes were acquired at 45 sec intervals. Acquisition settings were kept constant between groups: length of the movie, z-stack size and speed of acquisition. For the time course acquisition of T cell tracks and speed at 24h, 48h and 72h the same batch of animals has been imaged for 1 hour at the indicated time points.

### Movie analysis

T cell dynamics and T cell-tumor interactions were calculated using automated image analysis. For each movie, T cells and tumor cells were identified and tracked using the surface module of IMARIS software version 9.2.1, 9.3.1 and 9.5.1 (Bitplane). Cell positions were exported and processed using a custom developed Python extension. Data from appropriate movies were then aggregated and processed by a customized pipeline in Tibco Spotfire, and statistical analysis was calculated as stated in the figure legends via GraphPad Prism Version 6 and Version 7.

### Generation of humanized mice

NSG female mice (Taconic) aged 5 weeks were injected with Busulfan (15 mg/kg) followed by an injection of 100.000 human CD34^+^ cord blood cells (StemCellTechnologies) 24h later. 15 weeks after injection, humanized mice were screened for human T-cell frequencies by flow cytometry. Only humanized mice that revealed a humanization rate greater than 25% were used for efficacy studies.

### Efficacy study

1.5 *10^6^ WSU DLCL2 in 50% GFR Matrigel admixed with 50% RPMI were injected subcutaneously in HSC-NSG mice and the tumors were allowed to grow for approximately 20 days. Mice were assigned randomly into groups, by adopting a dedicated software. Only mice with a tumor volume between 200 and 800 mm^2^ have been included in the studies. CD20-TCB treatment was administered intravenously once per week at 0.5 mg/kg. Anti-human IFNγ (Bio X Cell, BE0235) or anti-human CXCL10 (ThermoFisher Scientific, MA5-23726S) was administered intravenously concomitantly with CD20-TCB in the designed groups. Tumor size was measured 3 times per week, scouts were taken 24 hours post second treatment. The subcutaneous tumors were resected and fixed in PFA 4% and later processed for FFPET (VIP6 AI, Sakura). 4 µm paraffin sections were subsequently cut in a microtome (Leica RM2235, Germany). Hematoxylin and Eosin staining was performed using standard protocol. Immunohistochemistry was performed in paraffin sections with the indicated anti-human antibodies: CD3 (SP7, Thermofisher); CD31 (DIA310, dianova); IFNγ (ab231036, Abcam); CXCL10 (LS-C312561, LS-Bio); CXCR3 (LS-B10183, LS-Bio); Ki67 (790-4286); DAPI (760-4196), by adopting the Leica autostainer platform following the manufacturer’s protocols (Leica Biosystems, Germany). CD3/CD31 duplex images were obtained with Iscan HT (Ventana) and analyzed with Definiens software for cell quantification. CXCR3, Ki67, IFNγ and CXCL10 images were obtained with a Vectra Polaris scanner and analyzed with Halo software for cell quantification.

### Statistical analysis

Statistical analysis was performed using Graph Pad Prism Version 6 and Version 7, and statistical tests were applied as stated in the figure legends.

## Results

### CD20-TCB-mediated killing requires the formation of stable T effector cell-tumor cell synapses

To characterize the cellular and molecular mechanisms of CD20-TCB-induced tumor cell killing, we analyzed the killing capacity of CD8^+^ T cells against two DLBCL cell lines: WSU DLCL2, and OCI-Ly18 **(Fig. 1a, Fig. supplementary 1a, 1b, 1f)**. We postulate that several intrinsic features of DLBCL models, including CD20 and ICAM-1 expression level, might modulate the sensitivity to treatment with CD20-TCB. Killing experiments aimed to correlate CD20 and ICAM expression with efficacy of killing in several DLBCL cell line showed that, there is no general correlation between those two factors when many cell lines are evaluated (data not shown), arguing in favor that there are several parameters involved in the tumor cell killing mediated by CD20-TCB. However, it is difficult to identify all factors that modulate killing in several lines of DLBCL by blocking several proteins involved in different step of the T cell killing pathway. In contrast, the formation of stable immune synapses between a human CD8^+^ T cell and a target cell is a fundamental step for T cell mediated cytotoxicity regardless of targeted tumor cells [23]. We thus hypothesized that the heterogeneity observed in CD20-TCB-mediated killing may rely on different functionality and stability of the formed immunological synapses. In order to investigate synapse formation *in vitro*, we took advantage of live confocal imaging and IMARIS built-in tracking algorithm to assess i) T cell dynamics (speed and movements direction); ii) T cells-tumor cells time of interaction and iii) area at the contact site over time. These three parameters help us defining “stable synapses”. We observed that CD20-TCB induces synapse formation between CD8^+^ T cells and both WSU DLCL2 or OCI-Ly18 cells **(Fig. 1b, supplementary Videos 1-4)** and, further supporting the hypothesis that these contacts mediate cytotoxic synapses, we observed polarization of the cytotoxic molecule Granzyme B toward the synapse **(Fig. 1c)**. Further analysis revealed that while T cells in untreated conditions are randomly moving, searching for a target, with a speed of ∼ 4 μm/min **(Fig. 1d, untreated)**, CD20-TCB treatment leads to a significant decrease in T cell speed **(Fig. 1d)**. Comparison of WSU DLCL2 and OCI-Ly18 cells revealed that the reduction in T cell speed is greater when CD8^+^ T cells target WSU DLCL2 cells, as compared to OCI-Ly18 cells **(Fig. 1d)**. In line with this, more T cells form interactions longer than 20 minutes when targeting WSU DLCL2 cells **(Fig. 1e)**. Furthermore, we observed that decreasing concentrations of CD20-TCB lead to less contact duration between T cell-target cells **(Fig. 1f)** and to a smaller area at the contact site **(Fig. 1g)**, supporting the role of CD20-TCB in tuning T cell-tumor cell synapse formation. Moreover, lower doses of CD20-TCB impair T cell speed, resulting in speed values similar to those found in untreated conditions **(Fig. 1h)**. Altogether, these data revealed that analysis of synapse formation between T cells and tumor cells, as a function of their motility and dynamic interactions *in vitro*, could be informative of T cells killing capacity upon CD20-TCB treatment. In support of the idea that stable synapses are required for efficient CD20-TCB-mediated tumor cell killing, we observed that the integrin protein with key functions in synapse formation and stabilization, LFA-1 [24], localizes at the synapse site **(Fig. 1i)**. Furthermore, while inhibition of FAS-L, an apoptosis trigger molecule, did not impact CD20-TCB mediated killing, LFA-1 inhibition decreased CD20-TCB mediated killing both in WSU DLCL2 cells **(Fig. 1j)** and in OCI-Ly18 cells **(Fig. supplementary 1c)**, without affecting T cells viability **(Fig. supplementary 1d)**. However, killing capacity at high dose of CD20-TCB (200 ng/ml) is unaffected by LFA-1 inhibition. Indeed, despite LFA-1 not localizing at the synapse site upon inhibition, T cells still present actin polarization and form synapses **(Fig. supplementary 1e)**, suggesting that a high dose of CD20-TCB can induce a stable synapse formation independently of LFA-1 engagement.

**Figure 1:**
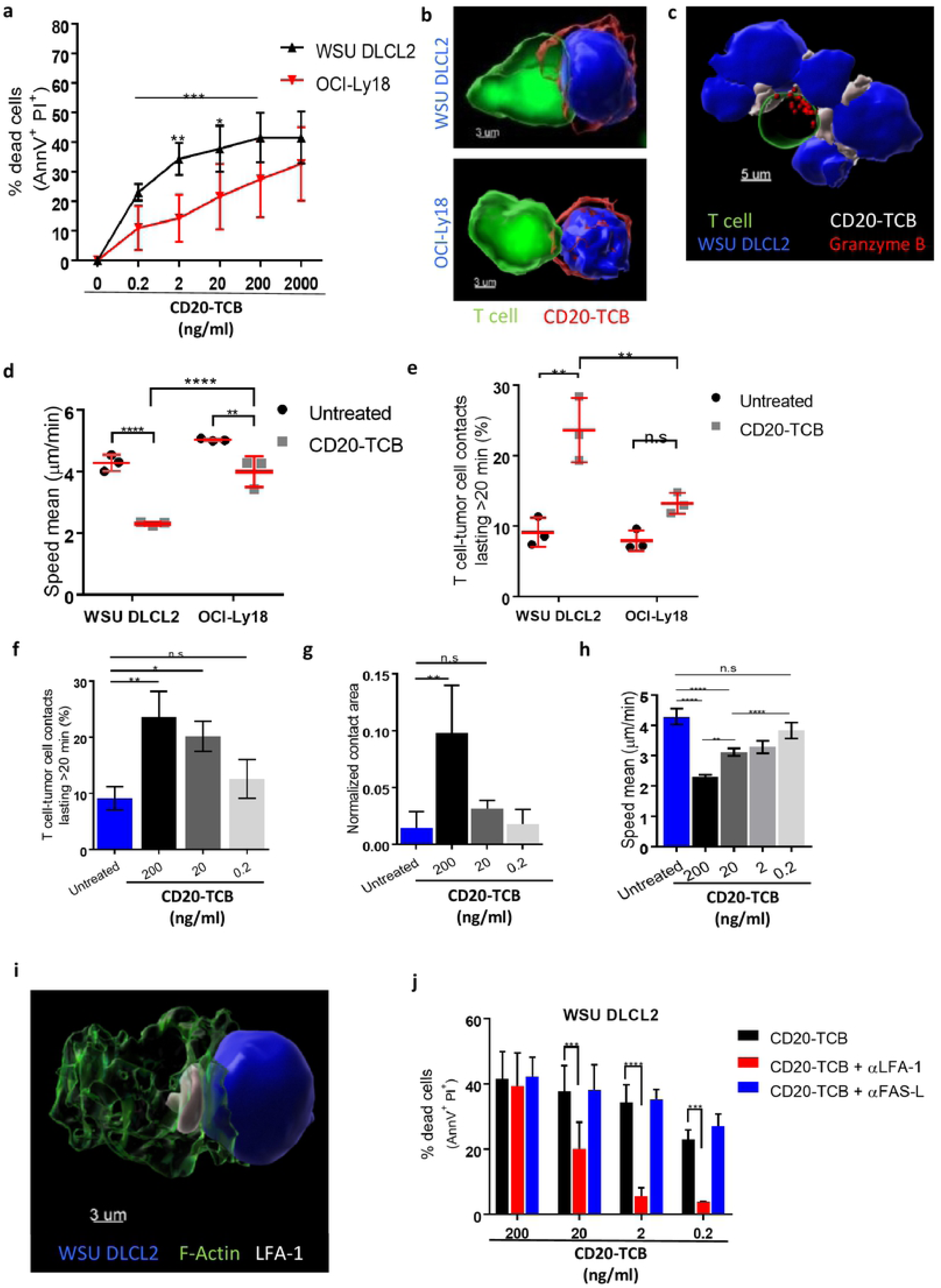
CD20-TCB mediates efficient tumor cell killing through stable synapses formation. **a)** Viability assay by AnnexinV (Annv^+^) and Propidium Iodide (PI^+^) staining on WSU DLCL2 and OCI-Ly18 cells, 16 hours after co-culture with CD3/CD28 activated CD8^+^ T cells in the presence of the indicated CD20-TCB concentrations. n=4 per point. Mean and +/- s.d. are shown. 2way-Anova, *p= 0.05, **p=0.005, *** p<0.0005. **b)** 3D reconstruction of representative confocal imaging of synapse formation between CD8^+^ T cell (green) and tumor cell (blue), mediated by CD20-TCB treatment (red). **c)** 3D reconstruct ion of representative confocal imaging of Granzyme B (red) polarization toward the synapse between CD8^+^ T cell (green) and tumor cells (blue), mediated by CD20-TCBtreatment (white). **d-e)** Confocal live cell imaging quantification of **(d)** Percentage of T cell-tumor cell contacts lasting more than 20 minutes; **(e)** CD8^+^ T cells speed (µm/min). Cells have been treated with +/- CD20-TCB (200 ng/ml), in the presence of WSU DLCL2 or OCI-Ly18 cells as target. Mean and +/- s.d. are shown, n=3. 2way-Anova, **p<0.005; ****p<0.0001; n.s.: not significant. **f-h)** Confocal live cell imaging quantification of **(f)** Percentage of T cell-WSU DLCL2 contacts lasting more than 20 minutes; **(g)** contact area between T cells and WSU DLCL2, normalized by the total surface of tumor cells and **(h)** CD8^+^ T cells speed (µm/min). Cells have been treated with CD20-TCB at the indicated doses. Mean and +/- s.d. are shown, n=3. 2way-Anova, *p<0.05; **p<0.005; ****p<0.0001; n.s.: not significant. i) Representative confocal imaging of LFA-1 (white) localization at the synapse between T cell (F-actin is shown in green) and target cell (blue) **j)** Viability assay by AnnV^+^ and Pl^+^ staining of WSU DLCL2 cells, 16 hours after co-culture with CD3/CD28 activated CD8^+^ T cells in the presence of the indicated CD20-TCB concentrations and +/- LFA-1 inhibitor (10 µg/ml) or +/- FAS-1 inhibitor (10 µg/ml). n=4 per point. Mean and +/- s.d. are shown. 2way -Anova, *** p<0.0005, **** p<0.0001.

These data revealed that CD20-TCB activity requires the formation of stable cytotoxic synapses and that the stability of these synapses correlates with the killing capacity. This parameter, which can be efficiently measured by live imaging approaches, is therefore key to explain the heterogeneity of the response of CD20-TCB against different cell lines of DLBCL. Despite peculiar features of target cells such as expression of CD20, ICAM-1 and LFA-1 might explain difference in killing between WSU and OCI-Ly18, they do not suffice to explain heterogeneity observed in other human DLBCL cell lines (**Fig. supplementary 1f and data not shown)**, leaving the stability of synapse formation the main read-out correlating with efficiency.

### A xenoreaction-free model for the visualization and quantification of infiltrating T cell dynamics in response to therapy

The selective binding of CD20-TCB to human CD20 on cancer cells, urged us to develop a suitable imaging model in order to visualize the response to CD20-TCB in the context of a human immune system *in vivo*. To this aim we developed an MP-IVM imaging experimental design in which we implanted a dorsal skinfold chamber in recipient NSG mice followed by intradermal injection of CMAC-Blue stained human DLBCL cells (WSU DLCL2) together with unstained murine CT26 cells, CMFDA-labeled human T cells and +/- AlexaFluor labeled CD20-TCB **(Fig. 2a)** [22]. To set up our MP-IVM studies we adopted two different sources of human T cells for the skinfold chamber of NSG mice, and we then compared the two humanized mouse models derived from these transfers. In one model (PBMC-NSG), we transferred labelled T cells derived from human peripheral blood mononuclear cells (PBMCs) in immune-deficient NSG mice. However, while this model is known to allow human T cell engraftment, it is prone to graft versus host disease (GvHD) due to strong xenoreaction of human T cells against the murine host tissue [25]. The second humanized model (HSC-NSG-NSG) was obtained by transferring in the skinfold chamber implanted in NSG mice human T cells derived from HSC-NSG mice. HSC-NSG mice were generated by transferring human hematopoietic stem cells (HSCs) from the cord blood of healthy donors into Busulfan pre-conditioned (and therefore depleted of endogenous bone marrow) NSG mice [26]. In this latter model, human T cells, after generation in the bone marrow, undergo selection in the murine thymus and are therefore tolerant to mouse tissue, thus minimizing xenoreaction. We adopted WSU DLCL2 cells for our *in vivo* imaging model, since *in vitro* experiments revealed that WSU DLCL2 targeting leads to more stable synapses as compared to OCI-Ly18 **(Fig. 1b-d)**, thus providing a powerful tool for investigating T cell dynamics *in vivo* in the context of CD20-TCB treatment. CT26 murine cells have been mainly adopted for technical reasons, as having WSU DLCL2 cells interspaced by CT26 cells allows for better segmentation of cells during post acquisition analysis, and therefore for a more precise quantification of T cell-target cell interactions. Moreover, the addition of CT26 cells allows the recreation of physiological heterogeneity within the implanted human tumor. Notably, CD20-TCB only targets WSU DLCL2 and not CT26 cells, and as CT26 cells are from a BALB/C background, thus sharing the expression of major histocompatibility complexes with NSG mice, tolerized T cells should not recognize them and cause alloreaction. Therefore, the introduction of CT26 cells in our model should not have an impact on tolerized T cells dynamics. We demonstrated that this approach allows for the simultaneous visualization, by MP-IVM, of tumor cells (blue), T cells (pink) and CD20-TCB (white) in three different channels **(Fig. supplementary 2a)**. In line with our *in vitro* observations, we could confirm that CD20-TCB binds concomitantly to tumor cells and T cells, forming visible clusters of increased drug density on the membrane of T cells **(Fig. 2b)**. Comparison of T cell dynamics revealed that T cells derived from PBMCs show a low motility both in the vehicle group **(supplementary Video 5)** and upon CD20-TCB treatment **(supplementary Video 6)**. In contrast, T cells from HSC-NSG mice were highly motile in the vehicle group **(supplementary Video 7)**, and slowed down visibly upon CD20-TCB treatment **(supplementary Video 8)**. To portray differences in T cell dynamics we displayed flower plots of tracks according to their displacement in the X and Y axes **(Fig. 2c)**. Tracks that go farther from the origin indicate lack of interaction with the surrounding tissue, while tracks that stay closer to the origin indicate a stronger interaction between T cells and the surrounding environment. The tracks of T cells in PBMC-NSG mice are not spreading far from the center, both in the CD20-TCB treated animals and in the vehicle **(Fig 2c, top)**, suggesting that in this model T cells interact with the tissue regardless of the treatment (xenoreactivity), thus providing an insufficient ability to evaluate differences in T cell dynamics upon therapy. On the contrary, T cells from HSC-NSG mice, when treated with CD20-TCB, displayed a track dispersion far lower than mice receiving only the vehicle **(Fig. 2c, bottom)**. In line with this, T cells in PBMC-NSG mice were almost immotile both in the untreated and CD20-TCB treated group **(Fig. 2d)**. In contrast, HSC-NSG derived T cells had an average speed of 6 µm/min in vehicle treated mice, which dropped to 3 µm/min upon CD20-TCB treatment **(Fig. 2d)**, recapitulating our *in vitro* observations **(Fig. 1g)**. Furthermore, T cells from PBMCs displayed no visible changes in track displacement from point A to point B in the acquisition frame, between untreated and treated groups **(Fig. 2d)**. By contrast, the track displacement of HSC-NSG-derived T cells was significantly different between the vehicle and the treated groups **(Fig. 2d)**, meaning that, without treatment, T cells are randomly moving and explore the environment far from their starting point, while CD20-TCB leads to T cells moving in a neighborhood closer to the starting point. These data revealed that by using our imaging experimental design we are able to visualize CD20-TCB impact on human T cell dynamics. Moreover, we were able to identify and describe the xenoreaction, which affects T cell dynamics in the intradermal region when utilizing human T cells originated from PBMCs. Further supporting the finding that PBMCs-derived T cells are immotile in the skinfold chamber due to ongoing xenoreaction with the host murine tissue, analysis of blood composition of PBMC-NSG mice revealed an increased proportion of strongly activated CD8^+^ T cells expressing Granzyme B and Ki67, a phenotype consistent with the ongoing GvHD **(Fig. supplementary 2b-c)**. By contrast, HSC-NSG mice are characterized by a higher % of circulating naïve T cells, defined as CD45RA^+^ CD62L^+^ **(Fig. supplementary 2d)**. These observations support the finding that T cells from HSC-NSG mice are tolerized in the murine thymus and therefore do not mediate xenoreaction.

**Figure 2:**
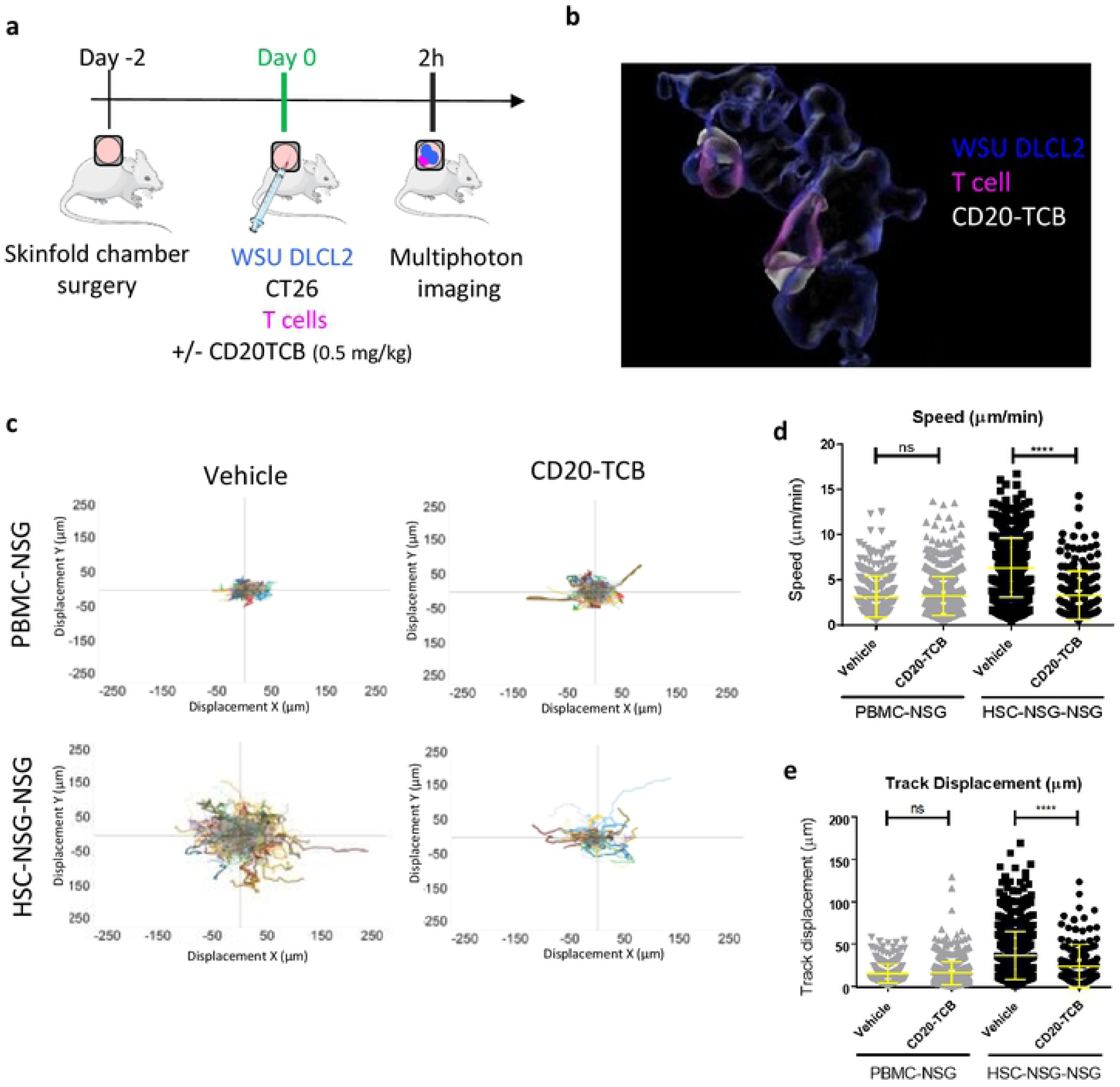
A xenoreaction-free model to quantify T cell dynamics in response to therapy. **a)** Workflow schematics: skinfold chambers were installed on NSG mice (day -2). 48 hours later (day 0): WSU DLCL2 (blue), unstained CT26 cells, and CD2^+^ T cells (pink) freshly purified from human PBMCs or from HSC-NSG mice were injected intra-dermally in the skinfold chamber together with labeled CD20-TCB (0.5 mg/kg) or with suitable vehicle. Cells were imaged 2 hours post treatment by MP-IVM. Adapted from https://smart.servier.com’. **b)** 3D representative rendering of MP-IVM imaging on skinfold chamber of HSC-NSG-NSG mice showing localization of therapy (white) at the contact site between WSU DLCL2 cells (blue) and T cells (pin k), 2 hours post treatment. **c)** MP-IVM analysis of T cells t racks in the skin fold chamber of PBMC-NSG (t op) *vs* HSC-NSG-NSG mice (bott om), +/- CD20-TCB.T cell tracks are plotted according to their displacement in the X and Y axes. Total number of tracks for each plot is: Top left: Vehicle n=330. Top right: CD20-TCB n=759. Bottom left: Vehicle n=741. Bottom right: CD20-TCB n=185. **d-e)** Quantification of **(d)** Track Speed (µm/min) and **(e)** Track displacement (µm) of T cells in PBMC-NSG or HSC-NSG-NSG mice, +/- CD20-TCB. Shown in yellow are mean values +/- s.d. Unpaired t-test; *** *p<0.0001;n.s.: not significant.

In conclusion, the model adopting HSC-NSG-mice-derived T cells constitutes the best approach to our aims as it minimizes xenoreaction and recapitulates CD20-TCB *in vitro* observations. Indeed, as observed *in vitro* **(Fig. 1g)**, CD20-TCB leads to decreased speed in T cell movements which is a consequence of the formation of functional synapses between T cells and target cells.

### CD20-TCB induces early and lasting T cell engagement and activity

Co-injection of CD20-TCB with tumor cells and T cells in the skinfold chamber revealed an early interaction of T cells with the tumor **(Fig. 2)**. However, this approach does not constitute a proper setting to describe CD20-TCB pharmacokinetics, as the drug is directly administered into the tumor not allowing the analysis of the impact of whole body distribution. To overcome this limitation, in a new set of experiments we administered CD20-TCB intravenously, aiming at understanding a) how long does it take for the drug to reach the tumor site and start inducing T cell-tumor cell interactions, as well as b) the duration of these interactions. In order to depict the degree of T cell-tumor cell interactions, we calculated different parameters from MP-IVM videos: T cell speed, track displacement from point A to point B in the acquisition frame, and arrest coefficient, which represents the percentage of T cells with a speed lower than the 2 μm/sec threshold. In order to understand the time point at which the CD20-TCB-induced stable contacts between T cells and tumor cells start to form, we determined an arbitrary threshold for “significant contact stability” by calculating the average values of the interaction parameters at 24h post therapy (a time point in which we know the interactions are stable upon treatment) **(Table 1)**.

**Table 1:** Reference average values of T cell dynamic indexes in tumors treated with 0.5 mg/kg CD20-TCB intravenously or with suitable vehicle, at early (less than 24 h) or late (more than 24h) time points. - The meandering index indicates the kind of movement of T cells. A meandering index of 0.5 indicates random movement, 1 indicates directional movement, and a meandering index of 0 indicates a cell that is completely engaged and almost doesn’t avert from its target.
- Track speed is an indication of the average speed of **T** cells, which decreases in the presence of interaction and engagement.
- Track displacement is another indicator of the way **T** cells move: the higher the track displacement, the further the cells migrate during acquisition, showing less interaction with the surrounding tumor.
- Contact duration represents the average contact duration between **T** cell tumor cells.
- The arrest coefficient indicates the percentage of cells that slow down to less than 2 µm/min, with a higher arrest coefficient meaning more arrested (and thus engaged) cells.

| Index | < 24 hours (early) |  |  |  |  |  | > 24 hours (late) |  |  |  |  |  |
| --- | --- | --- | --- | --- | --- | --- | --- | --- | --- | --- | --- | --- |
|  | vehicle |  |  | CD20-TCB |  |  | vehicle |  |  | CD20-TCB |  |  |
|  | Mean | S.E.M | N | Mean | S.E.M | N | Mean | S.E.M | N | Mean | S.E.M | N |
| Meandering index | 0.3783 | 0.006055 | 3 | 0.3085 | 0.01007 | 2 | 0.3968 | 0.006823 | 6 | 0.3369 | 0.005316 | 7 |
| Speed (µm/min) | 5.588 | 0.07982 | 3 | 2.448 | 0.104 | 2 | 4.42 | 0.07334 | 6 | 2.256 | 0.04495 | 7 |
| Track displacement (µm) | 32.28 | 0.6849 | 3 | 17.02 | 0.9613 | 2 | 23.3 | 0.5799 | 6 | 13.6 | 0.2876 | 7 |
| Contact duration (min) | 3.444 | 0.05564 | 3 | 7.74 | 0.2048 | 2 | 3.918 | 0.0758 | 6 | 7.343 | 0.1378 | 7 |
| Arrest coefficient (%) | 36.87 | 0.8229 | 3 | 72.7 | 1.39 | 2 | 44.06 | 0.9013 | 6 | 78.22 | 0.5674 | 7 |

Upon intravenous administration of CD20-TCB, the speed and track displacement of T cells decreased significantly and the arrest coefficient increased significantly, when compared to the vehicle, as early as 30 minutes after injection **(supplementary videos 9-14)**. However, the arbitrary threshold (shown as a dotted line), was reached at around 1-2 hours after therapy **(Fig. 3a)** for each of the parameters analyzed. These data suggest that CD20-TCB, despite being a large molecule (194.342 kDa), reaches the tumor site and acts on the resident T cells inducing T cell-tumor cell interaction early after systemic administration.

**Figure 3:**
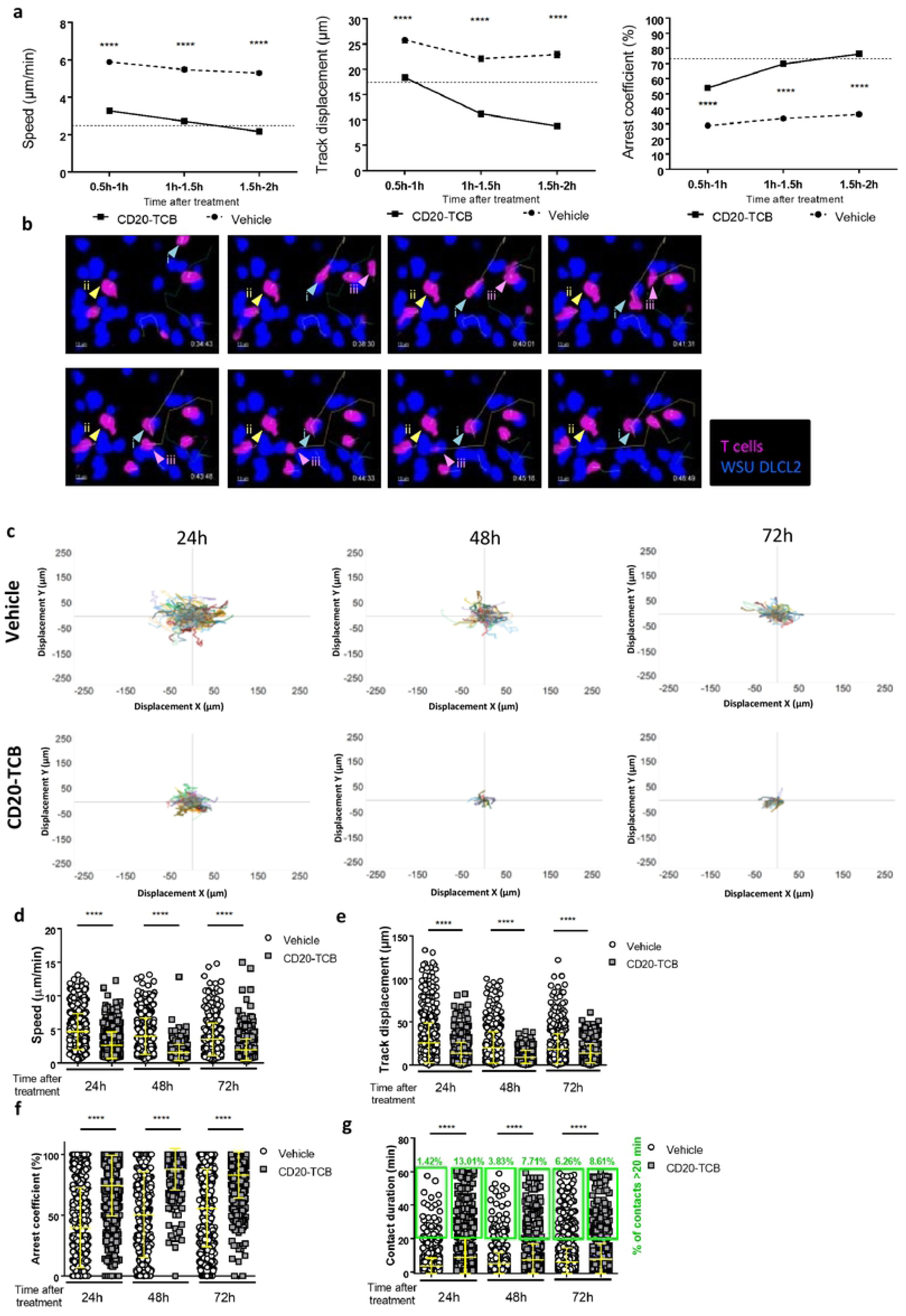
CD20-TCB induces early and lasting T cell engagement and activity. **a)** Time course quantification of Speed (µm/ min), Track displacement (µm) and Arrest coefficient(%) of HSC-NSG-derived T cells co-injected with WSU DLCL2 tumor cells in NSG mice, treated with 0.5mg/kg CD20-TCB i.v. or suitable vehicle and imaged starting from 30 minutes after treatment for 2 hours. Statistics: Kruskal-Wallis test, **** Adj. P value< 0.0001. Shown are mean values+/- SEM **b)** Time course tracking of T cells after intravenous injection of 0.5 mg/kg CD20-TCB. Highlighted are the 3 different possible behaviors of T cells: i) This T cell is moving fast at the beginning, with a straight trajectory. When it encounters a tumor cell in the presence of therapy, it suddenly stops, and starts interacting (light blue arrow). ii) This cell is interacting with the tumor since the beginning, it s track is revolving around the same coordinates (yellow arrow). iii) This cell does not interact with the environment (light blue arrow). WSU DLCL2 cells are represented in blue, T cell in pink. Tracks are shown for each T cell in the field. **c)** Shown are T cell tracks, plotted according to their displacement in the X and Y axes. Comparison of tracks of T cells treated intravenously with 0.5 mg/kg CD20-TCB or with suitable vehicle. Each animal has been imaged over time for 1h at 24,48 or 72h after treatment. 24h: Vehicle n=403, CD20-TCB n=348. 48h: Vehicle n=195, CD20-TCBn=49. 72h: Vehicle n=353, CD20TCB n =127. **d-g)** Quantification of **(d)** Speed (µm / min); **(e)** Track displacement (µm); **(f)** Arrest coefficient(%).**(g)** Contact duration (minutes) of T cells with tumor cells and percentage of contacts lasting more than 20 minutes (green boxes); T cells were derived from HSC-NSG spleens and co-injected with WSU DLCL2 tumor cells, treated with 0.5mg/kg CD20-TCB i.v. or suitable vehicle and imaged 24, 48 or 72 hours post treatment. Statistics: Kruskal-Wallis test, **** Adj. P value< 0.0001, ** Adj. P value< 0.005. Shown are scattered plots and means+/- s.d.

It is possible to visually appreciate the effect of CD20-TCB by observing 3 different behaviors of the T cells *in vivo*: i) a T cell migrating and suddenly recognizing a tumor cell and start interacting with the same; ii) a T cell interacting from the beginning of the observation with a tumor cell, and iii) a T cell that does not interact, possibly because it has not been engaged by CD20-TCB yet **(Supplementary video 15, Fig. 3b)**. Our observation confirms that CD20-TCB acts early upon injection (behavior ii), but also suggests a gradual effect of CD20-TCB on T cell dynamics (behavior i and possibly behavior iii).

To address the question of the duration of CD20-TCB–mediated T cell-tumor cell synapses, we investigated the interaction indexes at the tumor site at later time points. Flower plots of track displacement at 24, 48 and 72 hours consistently showed less dispersed tracks in the CD20-TCB treated tumors, suggesting a higher engagement of T cells at all time-points upon CD20-TCB treatment as compared to vehicle **(Fig. 3c and supplementary video 16-21)**. More in detail, analysis at 24, 48 and 72 hours post CD20-TCB injection revealed that both T cell speed and track displacement (distance covered by a single cell in a given time) decreased significantly when comparing vehicle *vs* treatment group. **(Fig. 3d-e)**. These data indicate that T cells form stable synapses with tumor cells also at later time-points upon CD20-TCB injection. The arrest coefficient values (percentage of cells moving slower than 2 μm/min at a given time point) also support the great stability and functionality of CD20-TCB induced synapses at all measured time-points. Indeed, in the treated group more than 70% of the cells were arrested at 24, 48 and 72 hours, while in the vehicle group always less than 56% of cells were arrested **(Fig.3f)**. Similarly, contact duration increased strongly upon treatment compared to the vehicle at all time-points, and the percentage of long term contacts (more than 20 minutes) increased remarkably when comparing vehicle *vs* CD20-TCB, with the biggest difference at 24 hours **(Fig. 3g)**. Overall, these data indicate a higher frequency and longer duration of T cell-tumor cell interactions in mice treated with CD20-TCB as compared to vehicle controls, not only at early, but also at late time points.

Altogether, these data show that CD20-TCB-dependent T cell-tumor cell interactions start gradually at early time points and persist at later time points. Moreover, the observation that CD20-TCB treatment leads to increased proportion of Granzyme B or Perforin expressing CD8^+^ T cells over time **(Fig. supplementary 3 a-b)** led us to conclude that these stable and persistent interactions are *bona fide* cytotoxic synapses, in line with those observed *in vitro* **(Fig. 1 c)**. However, besides CD20-TCB-mediated cytotoxicity, previous works had suggested that CD20-TCB could have additional activities within the tumor. Indeed, it has been observed that upon CD20-TCB treatment, an increased amount of intra-tumor T cells can be observed [3], possibly due to T cell influx form the periphery; however, the origin of this T cell influx has never been investigated.

### CD20-TCB treatment induces both the expansion of pre-existing intra-tumor resident T cells and the recruitment of peripheral blood T cells

We have demonstrated that CD20-TCB is able to mediate human T cell-tumor cell interactions in the presence of resident human T cells within 1 hour upon intravenous injection **(Fig 3)**. Moreover, in line with previous observations [3], we confirmed that CD20-TCB treatment leads to an increased number of CD3^+^ T cells in the tumor, as revealed by histology analysis of subcutaneously injected WSU DLCL2-derived tumors **(Fig. 4a)**. However, it is still not known whether the increased number of human CD3^+^ T cells could originate by resident T cell proliferation, recruitment from the periphery, or both. To this aim we collected tumors 24h post second injection of CD20-TCB or vehicle and performed histological quantification of intra-tumoral CD3^+^ T cells expressing i) Ki67, a proliferation marker; ii) CXCR3, the main receptor mediating recruitment of T cells to focal sites [27]. Our analysis revealed that CD20-TCB leads to increased counts of both CD3^+^ Ki67^+^ **(Fig 4b)** and CD3^+^ CXCR3^+^ **(Fig 4c)** cells, as compared to the vehicle, leading to the conclusion that CD20-TCB treatment leads to both proliferation of resident T cells and active recruitment of peripheral CXCR3^+^ cells. *In vitro* analysis revealed that CD20-TCB mediates early T cell activation, as demonstrated by CD69 and CD25 increased expression early upon treatment **(Fig. supplementary 4a)**, as well as T cell proliferation, **(Fig. 4d)**.

**Figure 4:**
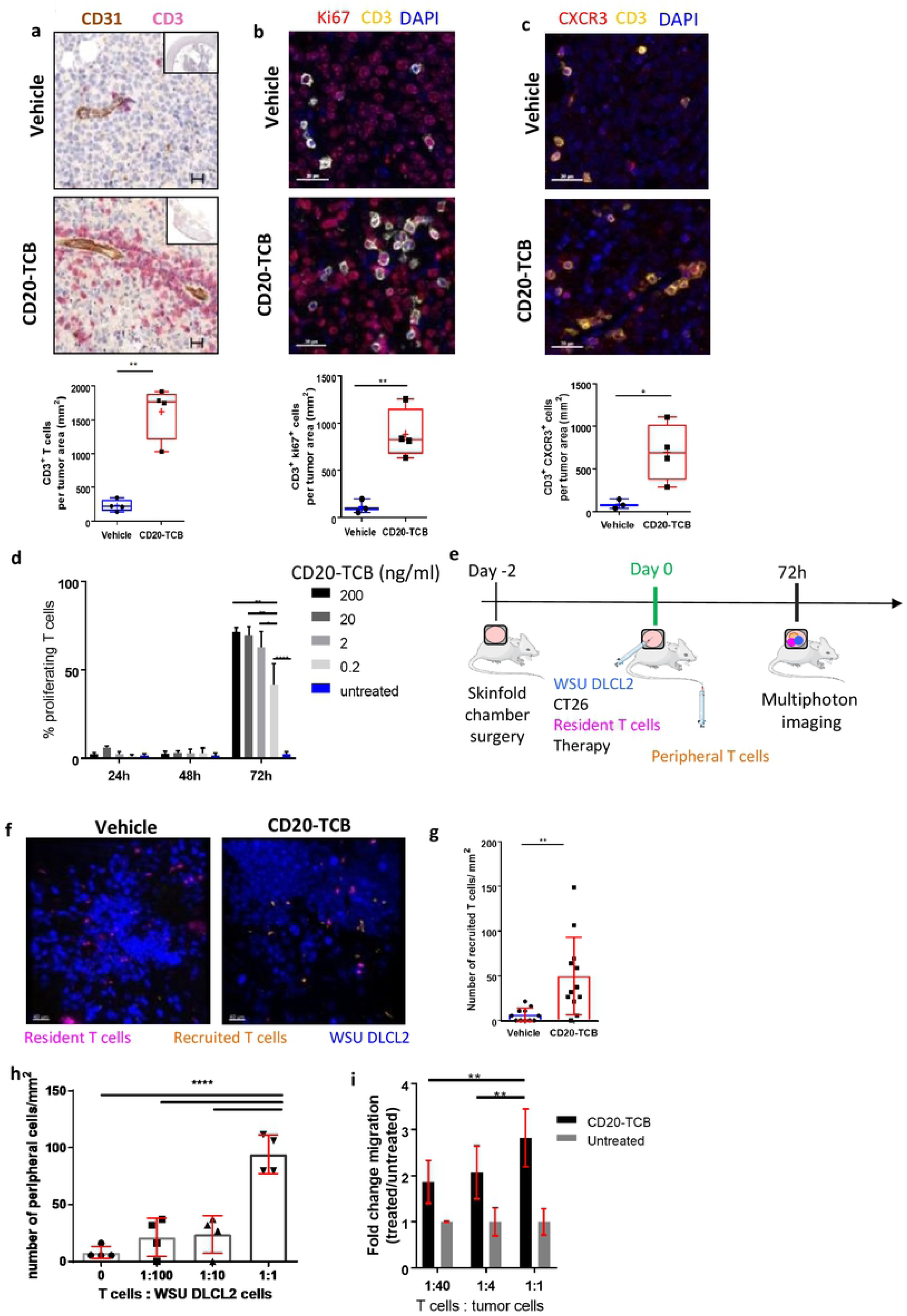
CD20-TCBinduces resident T cell proliferation and recruitment of peripheral blood T cells. **a-c)** Top: Representative histological staining of WSU DLCL2 tumors 24h post second treatment (0.5 mg/kg CD20-TCB or suitable vehicle i.v.). Bottom: Quantification of total number of cells/mm^2^ from histological images of vehicle vs CD20-TCB treatment. Whole slide scans quantification of 4 µm FFPE sections with the software **(a)** Definiens; **(b-c)** Halo. Statistical analysis: Unpaired 2-tailed t-test with Welch’s correction. *p<0.05, **p<0.005 **(a)** Red: CD3 staining, brown: CD31 staining. Quantification: number of CD3^+^ cells **b)** red: Ki67, yellow: CD3, blue: DAPI. Quantification: number of CD3^+^ Ki67^+^ cells **c)** Red: CXCR3, yellow : CD3, Blue: DAPI. Quantification of CD3^+^ CXCR3^+^ T cells. **d)** Percentage of proliferating CD8^+^ T cells, as assessed by CFSE dilution, freshly purified from PBMCs. Proliferation has been evaluated at 24h, 48h and 72h post CD20-TCB treatment, at the indicated doses, in the presence of WSU DLCL2 cells as target. n=3 per group, mean and s.d. are shown. One-way Anova, *p<0 .05, **p<0.005, ****p<0.0001 **e)** Workflow schematics: Skinfold chamber were installed on NSG m ice. 48h later, WSU DLCL2 (Blue), unstained CT26 cells, and CD2^+^ T cells freshly purified from HSC-NSG spleens (pink) were injected intra-dermally in the skinfold chamber, together with 0.25 mg/kg of CD20-TCB or with suitable vehicle. Concomitantly, freshly purified CD2^+^ T cells from HSC-NSG spleens (orange) were injected i.v. to allow visualization of peripheral blood T cells. Cells were imaged 72h post treatment by MP-IVM. **f)** Representative MP-IVM imaging of the tumors. Blue: WSU DLCL2 cells; Pink: Resident T cells; Orange: Recruited T cell s. Images were acquired 72h post intradermal treatment with 0.25 mg/kg CD20-TCB or suitable vehicle. Adapted from https://smart.servier.com/ **g)** Quantification of peripheral T cells (number/ mm ^2^) 72h post treatment. Mean +/- s.d. are shown. Unpaired 2-tailed t-test with Welch’ s correction. * * p<0.005. **h)** In the context of the skinfold chamber model, increasing number of T cells (Resident) were co-injected with the tumor and 0.25 mg/ kg of CD20-TCB intradermally, while 2.5*10^6^ T cells were injected intravenously (Peripheral). 72h post treatment, peripheral blood T cells were counted for each tumor from 5 representative fields. 4 tumors per group were analyzed. Shown is the count of peripheral T cells/ mm^2^, Mean+/- s.d. per group. Statistical analysis: One-way Anova. **** p<0.0001. **i)**3 hours *in vitro* chemotaxis assay of T cells toward preconditioned medium derived from WSU DLCL2 co-culture with CD3/CD28 pre -activated T cells. Pre-activated CD8 T cells have been plated with WSU DLCL2 cells at decreasing T cells : tumor cells ratios, in the presence of 200 ng/ml of CD20-TCB.24h later the supernatant has been collected and transferred to the bottom chamber of a 24-Transwe ll plate. In the top chamber 100.000 pre activated T cells, labeled with CFSE, have been seeded and let to migrate for 3 hours. Migration has been evaluated by counting total amount of CFSE positive migrated cells in the bottom chamber, by flow cytometry at constant volume and acquisition speed. Mean fold change and +/- s.d. are shown. n=5, from two independent experiments 2-way Anova; **p<0.005

In order to investigate the mechanism by which CD20-TCB activity within the tumor mediates peripheral human T cell recruitment *in vivo*, we adapted our skinfold chamber model. As the focus of the study are the mechanisms of active recruitment mediated by the activity of the drug within the tumor, we injected labelled resident T cells together with the tumor and the TCB intradermally, in order to provide a model of an “inflamed” tumor in which the drug is already active. At the same time, we injected differently labelled T cells intravenously. In the presence of any recruiting mechanism dependent on the CD20-TCB activity within the tumor, the peripheral T cells would be recruited to the tumor site, and would be easily distinguishable from their resident counterpart **(Fig. 4e)**. Our results clearly demonstrated that treatment with CD20-TCB strongly increases the intra-tumor infiltration of peripheral blood T cells (orange) as compared to the vehicle **(Fig. 4f)**, increasing the amount of T cells/mm^2^ from an average of 6 cells/mm^2^ in the vehicle to 50 cells/mm^2^ in the CD20-TCB treated group **(Fig. 4g)**.

In line with the observation that intravenous injection of CD20-TCB leads to early and lasting activity, intradermal CD20-TCB retains the ability to stimulate the interaction between resident T cells and tumor cells at 72 hours post treatment. Indeed, also in this setting, T cells interacted more with the tumor in the presence of CD20-TCB, as shown by the decrease in the speed index from an average of 5 µm/min in the vehicle to an average of 2.5 µm/min in the treated group. Likewise, track displacement and the arrest coefficient indexes indicated a higher T cell-tumor cell interaction in the treated *vs* untreated group. Moreover, the recruited T cells showed interaction parameters similar to the resident T cells in the treated group, supporting the finding that CD20-TCB can still bind freshly recruited T cells and is functional at 72 hours post administration **(Fig. supplementary 4b)**.

An important question to answer, is whether the presence of resident intra-tumor T cells at baseline is required to induce further T cell influx upon CD20-TCB treatment, and most importantly, whether there is a threshold of baseline infiltration required to mediate this secondary recruitment effect. To address these questions, we injected decreasing T cell-tumor cell ratios intradermally, and quantified the ability to recruit peripheral T cells upon CD20-TCB treatment. A 1:1 proportion of T cells *vs* tumor cells allowed to recruit a strikingly high number of peripheral blood T cells (up to 111 cells per tumor, with a minimum of 79 cells per tumor); decreasing the ratio to 1:10 or 1:100 strongly decreased the recruitment capability, although this was still retained (up to 37 cells per tumor). Notably, with no resident human T cells injected, the number of peripheral human T cells found in the tumor was extremely low (maximum 15 cells per tumor), similarly to the number of peripheral T cells detected in the vehicle group. These findings strongly suggest that the presence of a certain amount of resident T cells is necessary to trigger the initial inflammation and T cell proliferation as well as to induce peripheral T cell recruitment upon TCB treatment **(Fig. 4h)**, hence transforming a desert tumor, with limited number of resident T cells, in an inflamed tumor.

By *in vitro* chemotaxis assays, we showed that supernatants derived from WSU DLCL2 co-cultured with CD8^+^ T cells and treated with CD20-TCB lead to peripheral human T cells recruitment (from the top chamber) **(Fig. 4i, ratio 1:1)**. Moreover, we observed that supernatants derived from lower T cell-tumor cells ratios have a lower attractant capacity **(Fig. 4i)**. Altogether, these data suggest an active recruitment activity exerted by resident T cells when exposed to CD20-TCB. However, the mechanism through which CD20-TCB leads to the recruitment of peripheral human T cells still remains to be elucidated.

### CD20-TCB-induced peripheral T cell recruitment is dependent on the IFNγ-CXCL10 axis

Based on the finding that CD20-TCB mediates the recruitment of peripheral human T cells into the tumor, we hypothesized that this might be mediated by chemokines released after T cell-tumor cell contact. It has long been known that cytotoxic T cells produce, among many other cytokines, also high level of IFNγ [28, 29]. One of the most common recruitment axes for peripheral T cells into tumor involves this cytokine [30]. Quantification of chemokines levels in the supernatant from PBMCs derived CD8+ T cells co-cultured with WSU DLCL2 cells, revealed that, among other chemokines and cytokines, IFNγ increased over time after CD20-TCB treatment, as compared to the untreated condition, reaching the peak of INFγ levels 72h post treatment **(Fig. 5a)**. In the same settings, we indeed observed an increased proportion of IFNγ^+^ CD8^+^ T, as compared to the untreated condition, starting from 48h post treatment **(Fig. 5b)**. These data support the hypothesis that CD20-TCB first leads to stable synapses formation, that mediate direct tumor cytotoxicity and cytokine and chemokines production which leads to additional effects of CD20-TCB. Histology analysis of IFNγ levels in subcutaneous tumors, despite heterogeneity of the samples, revealed a trend (p=0.057) in higher amount of IFNγ levels, of around 30 fold, 24h post second injection of CD20-TCB, as compared to the untreated group **(Fig. Supplementary 5a)**. It is well established that local IFNγ production can mediate the recruitment of CXCR3^+^ effector T cells to tumors [31]. The main chemokines involved in CXCR3^+^ T cell recruitment to tumors are CXCL9, CXCL10 and CXCL11 [32]. In order to assess whether IFNγ sustains CXCL9, CXCL10 and CXCL11 production in our settings, we treated WSU DLCL2 cells with decreasing doses of recombinant IFNγ *in vitro* and then measured the abundance of these chemokines in the supernatant. Our analysis revealed that IFNγ induces a selective and significant expression of CXCL10 in tumour cells**(Fig. 5c)**. In line with this, CD20-TCB treatment of human T cell-WSU DLCL2 co-cultures leads to increased levels of many cytokines and chemokines including CXCL10, most likely by both tumor cells and T cells **(Fig. 5d and data not shown)**. Based on the increase in CXCL10 expression upon IFNγ or CD20-TCB treatment that we have observed, we then tested CXCL10 amount in the supernatant obtained from the co-culture of human PBMCs-derived CD8^+^ T cells with WSU DLCL2 cells at 24h, 48h and 72h post CD20-TCB treatment. Our analysis revealed that CD20-TCB leads to increased CXCL10 secretion overtime that reaches the peak at 72h post treatment, as previously observed for IFNγ **(Fig. 5e)**, strongly suggesting a direct correlation between the two mechanisms. In addition, tumor tissue analysis by histology revealed the same trend of increased CXCL10 expression **(Fig. supplementary 5b)** in mice treated with CD20-TCB as compared to vehicle (increase of about 22 fold). In order to prove the role of IFNγ in the recruitment of peripheral T cells, we pre-treated WSU DLCL2 with IFNγ before intradermal injection, and imaged the tumors 72 hours later **(supplementary video 24)**. Our analysis revealed an increased recruitment of peripheral blood T cells (orange) by IFNγ pre-treated tumors, as compared to CD20-TCB **(supplementary video 23)** or vehicle **(supplementary video 22)** treated tumors. Moreover, analysis of T cell dynamics rules out the hypothesis that the recruitment is due to an increased activity of the resident T cells (pink) as a consequence of IFNγ pre-treatment **(Fig. Supplementary 5c)**. This finding suggests that CD20-TCB-mediated peripheral T cell recruitment *in vivo* might indeed involve factors released by tumor cells stimulated by T cell-produced IFNγ.

**Fig 5:**
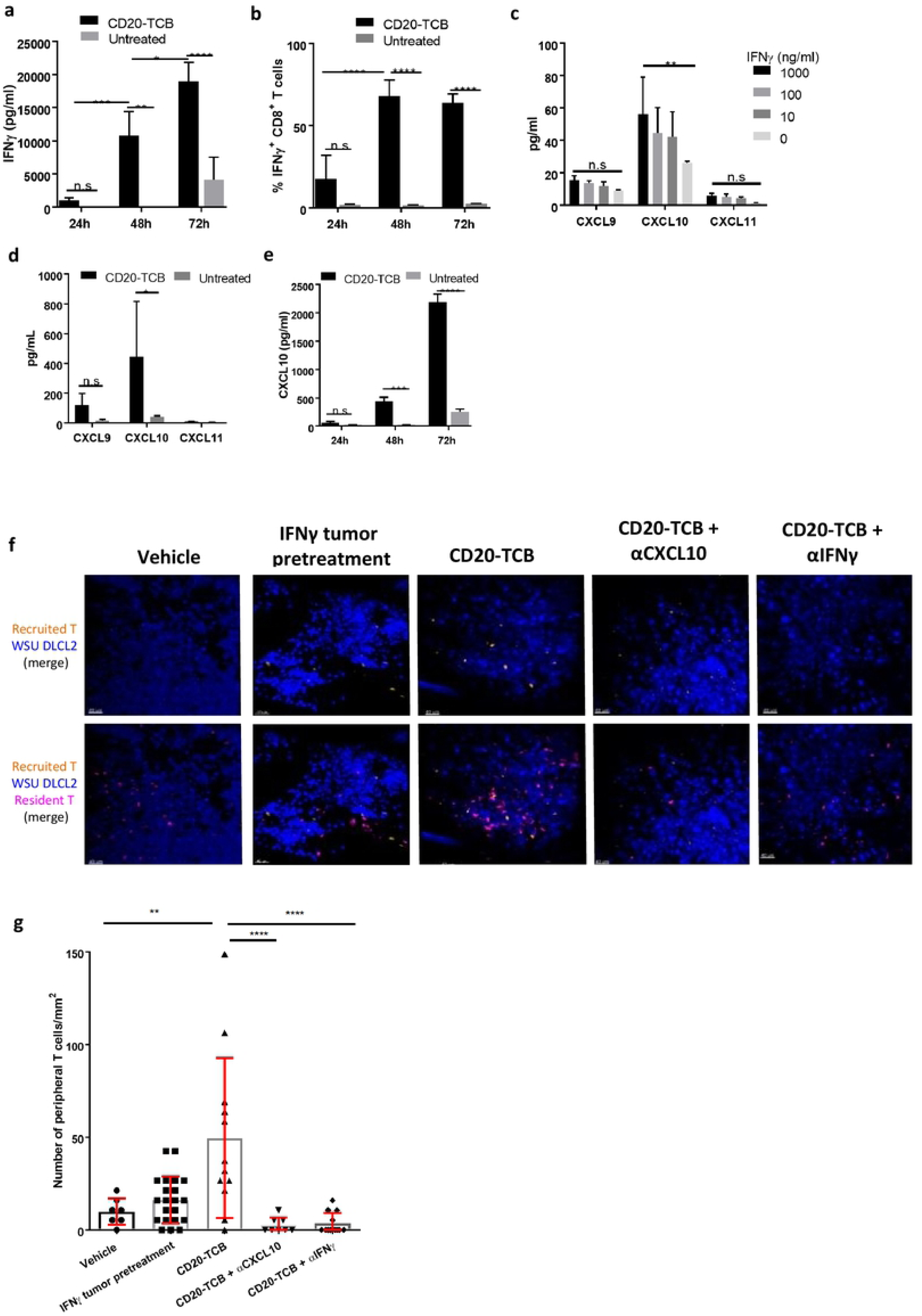
CD20-TCB-induced T cell recruitment is dependent on IFN γ and CXCL10. **a)** IFNγ protein quantification by multiplex analysis of supernatant derived from co-culture of WSU DLCL2 cells with CD8^+^ T cells freshly purified from PBMCs and stimulated with C0 20 -TCB(200 ng/ml) at the indicated time points. n=3 per group. Two-way Anova. *p< 0.05, **p< 0.05, ***p < 0.001, ****p < 0.0001, n.s.: not significant. **b)** Flow cytometry analysis of CD8^+^ INFγ^+^ T cells, at 24, 48 and 72 hours post *in vitro* CD20-TCB (200 ng/ml) treatment. **c)** Quantification of released cytokines (pg/ml) upon IFNγ stimulation of WSU DLCL2 tumor cells for 48 hours. Shown is Mean +/- s.d. of 3 replicates. **d)** Quantification of released cytokines (pg/ml) upon CD20-TCB treatment (200 ng/ml) of WSU DLCL2 co-cultured with CD3/C028 pre-activated CD8^+^ T cells. Shown is Mean +/- s.d. n=4 per group. Two-way Anova *p< 0.05, n.s.: not significant. **e)** CXCL10 protein quantification by multiplex analysis of supernant derived from co-culture of WSU DLCL2 cells with CD8^+^ T cells freshly purified from PBMCs and stimulated with C0 20-TCB (200 ng/ml) at the indicated time points. n=3 per group. Two-way Anova. ***p <0.001, ****p < 0.0001, n.s. not significant. **f)** Representative images from MP-IVM imaging in the skinfold chamber of HSC-NSG-NSG mice. WSU OlCL2 cells (blue) pre-treated or not with IFNγ were injected intra-dermally together with CD2^+^ T cells derived from the spleen of HSC-NSG(pink) and 0.25 mg/kg C0 20 -TCB or suit able vehicle. CD2^+^ T cells derived from the spleen of HSC-NSG (orange) where concomitantly injected intravenously. Top row: Tumor cells (blue) and peripheral T cells (orange). Bottom row: tumor cells (blue), resident T cells (pink) and peripheral T cells (orange). Where indicated, antibodies against CXCL10 or IFNγ were injected intravenously.**g)** Quantification of peripheral blood T cells (count/ mm^2^) 72 hours post treatment. Shown are individual counts/ mm^2^ and mean +/- s.d. Statistical analysis: One-way Anova. ***p <0.001,****p <0.0001.

To prove the role of IFNγ and CXCL10 in T cell recruitment, we performed an experiment comparing skinfold chamber tumors pre-treated or not with IFNγ, tumors treated with CD20-TCB, and tumors in which the treatment with CD20-TCB was followed by intravenous injection of αIFNγ or αCXCL10 blocking antibodies. We then imaged the tumors 72 hours post therapy **(Fig. 5f)**, and we counted the peripheral blood T cells (orange) recruited into each tumor **(Fig. 5g)**. In tumors that had been previously treated with IFNγ, we could see a tendency of an increase in the number of peripheral blood T cells with a maximum of 42 peripheral blood T cells per mm^2^, although the recruitment was significantly higher in CD20-TCB-treated tumors, with a maximum of 148 peripheral blood T cells per mm^2^. The presence of either blocking antibody against CXCL10 or IFNγ was able to significantly reduce peripheral T cell recruitment to a maximum of 5-10 peripheral blood T cells per mm^2^ **(Fig. 5g)**. This finding supports the hypothesis that, upon treatment with CD20-TCB, resident T cells start killing the tumor, increasing the inflammatory cytokines, such as IFNγ, in the microenvironment. This in turn induces tumor cells and T cells to secrete CXCL10, mediating the recruitment of CXCR3^+^ T cells from the periphery. As previously stated, the amount of tumor resident T cells impacts on the capacity to induce peripheral T cell recruitment upon treatment **(Fig. 4h-i)**. In line with this we observed that decreasing the number of T cells in co-culture with WSU DLCL2 cells, leads to decreased amount of IFNγ and CXCL10 secretion **(Fig. supplementary 5d-e)**. To further validate the importance of this part of CD20-TCB MoA, we performed an *in vivo* efficacy experiment by subcutaneously injecting WSU DLCL2 cells, and treating established tumors with vehicle, CD20-TCB, or CD20-TCB in combination with αCXCL10 or αIFNγ blocking antibody **(Fig. 6a)**. All treated groups were able to mediate tumor growth control and tumor regression, with CD20-TCB alone being able to almost eradicate the tumor by day 38 post therapy. The blocking of CXCL10 and IFNγ was able to slightly reduce the anti-tumor effect of CD20-TCB, suggesting that their inhibition might be able to at least partially block T cell recruitment to the tumor. This was confirmed by histology examination whereby tumors treated with CD20-TCB showed a homogeneous and strong infiltration of CD3^+^ T cells, while treatment with either αCXCL10 or αIFNγ was able to partially reduce intra-tumor T cell infiltration **(Fig. 6b, 6c)**. The tumors in the different groups displayed a high variability in size and infiltration; however, when investigating the correlation between tumor infiltration and tumor volume at termination, a significant inverse correlation was observed, with higher infiltration correlating with a stronger efficacy of CD20-TCB, thus a smaller tumor volume **(Fig. 6d)**. Overall, these data show that CD20-TCB treatment is efficacious in mediating tumor infiltration and subsequent cytotoxicity. Moreover, we identified the IFNγ-CXCL10 axis as being one of the drivers of TCB-mediated T cell infiltration in tumors, whereby IFNγ released by activated T cells induces CXCL10 secretion, which in turns attracts CXCR3-expressing T cells into the tumor sites. Furthermore, we could show a strong correlation between infiltration and tumor shrinkage, regardless of the therapy regimen.

**Fig 6:**
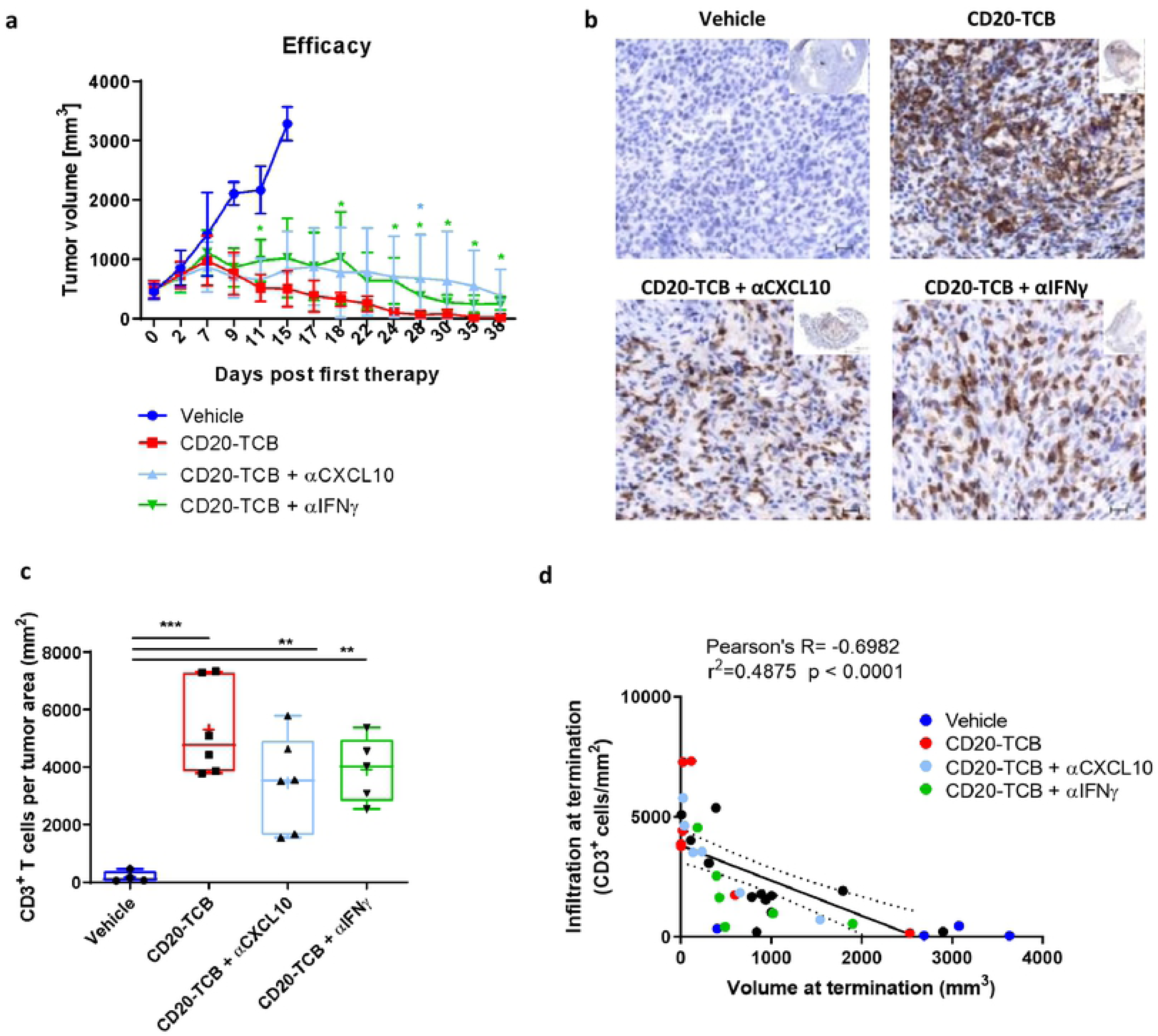
Inhibition of CXCL10 or IFNγ impairs CD20-TCB-dependent T cell infiltration. **a)** Subcutaneous WSU DLCL2-derived tumors between 200 and 800 mm^3^ were intravenously treated with 0.5 mg/kg CD20-TCB in the presence of aCXCL10 or αlFNγ antibodies (10 mg/kg), or with suitable vehicle, once per week. n=9. Shown are mean values +/- s.d. for tumor volume over time. Unpaired t test, *p<0.05. **b)** Representative histology of tumors at termination. Blue: nuclear counterstain, brown: CD3. **c)** Quantification of infiltration at termination (CD3^+^ T cells per tumor area). Statistics: One way Anova, **p< 0.01, ***p<0.005. **d)** Correlation between infiltration and tumor growth.

## Discussion

Redirecting T cells to hematological malignancies using bispecific antibodies is an attractive strategy to improve clinical outcome in addition to standard of care treatments. CD20-TCB (also known as RG6026 or glofitamab) is one of such examples showing remarkable benefits for relapsed and refractory DLBCL patients [4, 12, 13, 33]. Like other bispecific antibodies, CD20-TCB activates T cells by selectively crosslinking T cells to tumor cells through the simultaneous binding of the CD3ε invariant TCR signaling component and a tumor antigen (TA), specifically CD20 [34]. Using the aggressive DLBCL xenograft model (WSU DLCL2) and preclinical animal models of stem cell humanized (HSC) NSG mice[33], we have previously reported the mechanisms leading to CD20-TCB mediated *in vitro* tumor cell killing and its efficacy *in vivo* [3]. In this work, we had found that CD20-TCB can mediate cross-linking of T cells to tumor cells which correlated with CD20-TCB localization at the synaptic area [3], but information on the impact of the TCB mediated synapse quality (speed, contact length, area of contact, etc…) on the functional outcome (killing, recruitment, cytokine release, etc…) were still missing. One hypothesis arising from those findings was that CD20-TCB MoA, similar to TCR mediated interaction, could facilitate the formation of stable immunological synapses between T cells and tumor cells, whose stability and quality would impact T cell activation, cytotoxicity and cytokine release, as demonstrated in the context of TCR-mediated immune synapses [35]. To address this, we took advantage of the lower efficacy of CD20-TCB to induce T cell killing of OCI-Ly18 target tumor cells (as compared to WSU DLCL2). We realized that lower killing ability correlated with differential motility of T cells in culture with tumor cells. T cells in contact with OCI-Ly18 showed a lower contact area with tumor cells and increased speed as compared to T cells cultured with WSU DLCL2. These data supported our hypothesis that CD20-TCB affects the formation of T cells-tumor cells synapses. One of the major components of the immune synapsis is the integrin LFA-1 [15], which has been shown to be upregulated on canonical T cells surface after CD20-TCB crosslinking [3]. Among the multiple factors involved in synapse formation, the integrin LFA-1 is required for the stabilization and structural organization of the immunological synapse, by binding to its ligand ICAM-1 on target cells [36]. By directly visualizing LFA-1 using confocal microscopy, we observed that LFA-1 localizes at the synapse site between tumor and T cells. In addition, by selectively blocking LFA-1, we impaired the killing capacity of T cells stimulated by low-dose (but not high dose) of CD20-TCB. Because T cell killing was not impaired by blocking canonical apoptosis ligands such as FAS-L, we concluded that CD20-TCB-induced synapse formation is very important for its activity [37, 38]. This is not surprising as the synapse area is a critical parameter for effector T cells activity, including release of effector cytokines, effector granules and death inducing ligands [39]. In addition, inhibitory pathways such as PD-1 and CTLA-4 are recruited at the synapse area and may interfere with synapse formation [40, 41]. Altogether, our data suggest that CD20-TCB improves T cell killing of tumor cells by stabilizing synapses. T cells-tumor cells synapse stabilization may also interfere with negative feedback loops and thereby CD20-TCB could have synergistic effects with checkpoint inhibitors in the context of cancer immuno-therapy. The efficacy of CD20-TCB combination with PD-L1 inhibition by Atezolizumab is currently being investigated in a Phase Ib study (NCT03533283).

Given the importance of the synapse formation by CD20-TCB to mediate activity, we explored innovative models to perform dynamic *in vivo* imaging [22, 42]. Indeed, evaluation of synapse formation between human T cells and tumor *in vivo* using clinical lead molecules has been a long lasting question that was never optimally addressed due to lack of a proper model. One of the main reasons of failure is due to the xenoreaction of human cells towards mouse tissues and to the alloreaction of T cells towards the tumor [18, 25, 26]. These two reactions can strongly affect the T cell behavior possibly masking any additional effect of the studied therapy. We hereby present an experimental design that aims at reducing xenoreaction thus allowing the study of the T cell-tumor cell dynamics in response to immunotherapy. We show here that GvHD does not occur in stem cell humanized mice (HSC-NSG mice), as opposed to human PBMC transfer into NSG mice (PBMC-NSG mice). The profile of leukocytes derived from HSC-NSG mice more closely resembles the profile of human blood-derived cells than the profile of leukocytes derived from mice recipients of PBMC transfer, with lower frequency of activated and higher frequency of naïve conventional T cells. Furthermore, T cells derived from the spleen of HSC-NSG humanized mice where less reactive to the tumor environment in the absence of therapy when compared to T cells derived from human PBMCs. It was indeed possible to observe a difference between treated and untreated tumors in terms of T cell response, indicating that the transfer of humanized mice-derived T cells in the context of NSG mice (HSC-NSG-NSG) is a good model to analyze T cell dynamics in tumors, minimizing xenoreaction. The control group for the alloreaction towards human tumors that might still occur, was a vehicle group injected with the same T cell batch. One limitation of this model, as compared to syngeneic models [43] or other humanized mouse models [25, 26], is that the most represented functional cells are conventional CD4 and CD8 T cells [3] and therefore the conclusion on the mode of action of CD20-TCB mostly refers to those subsets. Treg CD4^+^ cells which are very important in controlling tumor growth and response to immunotherapies [44] are poorly represented. To verify whether these findings also affect Treg, the platform here described could be adapted using humanized models with higher degree of Treg frequency [45].

Our innovative model allowed us to evaluate *in vivo* the direct effect of CD20-TCB on T cell dynamics and synapse formation in the tumor upon intravenous administration, correlating pharmacokinetics with the pharmacodynamic changes in the tumor defined as synapse formation between human T cells and tumor target cells. Despite CD20-TCB high molecular weight (194.342 kDa), we found that T cells start interacting and forming synapses with the tumor cells as early as 30 minutes after intravenous injection of CD20-TCB. In addition, the analysis of the dynamics of recruited T cells at 72 hours after therapy, indicate that CD20-TCB is still available in the tumor and it can still mediate interaction between tumor and T cells. We here presented a novel imaging platform, flexible enough to be applied to analyze additional leukocyte subpopulations in the context of T cell bi-specific engagers such as TCB. This could help define the time a given compound reaches the tumor site and activates a response, making it possible to evaluate functional pharmacokinetics and the duration of the effect of the drug after treatment, possibly applying this knowledge to the design of treatment schedules for preclinical and clinical studies. Besides its cross-linking and direct cytotoxicity effect, it has been observed that CD20-TCB, as well as other T cell bispecific antibodies, lead to an increased number of intra-tumor CD3 T cells when compared to vehicle, probably as a result of CD20-TCB-induced T cell proliferation and cytokine production [3, 43, 46]. By histological analysis of tumors treated with CD20-TCB, we revealed a significant increase of intra-tumor proliferating CD3 T cells, as compared to the vehicle. In addition, we observed increased numbers of CXCR3^+^ CD3^+^ T cells within the tumor, suggesting a possible CD20-TCB-dependent recruitment of peripheral activated and memory T cells, in line with previous observation using TCR mediated T cell activation [47, 48]. A previous study in a breast cancer mouse model, using a surrogate molecule, showed that HER2-TDB leads to increased numbers of CXCR3^+^ CD3^+^ murine T cells within the tumor [43], suggesting that this mechanism might be shared among different T cell engagers. However, it still remains to be addressed whether human T cells share a similar behavior with murine T cells. Our data suggest indeed that also human T cells use the axis IFNg-CXCL10-CXCR3 as one of recruitment mechanism of peripheral T cell into tumor.

An important observation supporting the elucidation of recruitment of human T cells into the tumor is that the amount of peripheral T cells recruited to the tumor correlates with the number of resident T cells. Indeed, 72 hours post therapy, the absence of resident T cells in the tumor had no effect on peripheral T cell recruitment, resulting in comparable effects to those observed in vehicle. By contrast, even a low starting ratio of T cells-tumor cells (1:100 and 1:10) was sufficient to mediate a detectable infiltration of peripheral blood T cells when compared to vehicle. Noteworthy, increasing the number of resident T cells to a 1:1 ratio to tumor cells further increased the number of recruited peripheral blood T cells. This finding supports the idea that CD20-TCB ability to engage the few resident T cells present in tumor is important to mount the intra-tumor inflammation and lead to anti-tumor efficacy and plays an important role in transforming desert tumor to tumor infiltrated by T cells. [43, 46, 49].

As previously shown, some of the chemokines involved in T cell recruitment to tumors are the Chemokine (C-X-C motif) Ligands (CXCL)-9 and 10 [32]. These are induced by local TNFα and IFNγ cytokine production, and can mediate the recruitment of Chemokine Receptor (CXCR)-3-positive effector T cells to tumors [31]. Different studies demonstrated that CXCR3^+^ T cell recruitment is regulated by IFNγ [50]. In our study we demonstrated that CD20-TCB leads to increased IFNγ secretion by activated T cells. Among the CXCR3 binding chemokines produced, we focused on the increase of IFNγ and CXCL10, after CD20-TCB treatment. By using blocking antibodies *in vivo*, we confirmed that the IFNγ-CXCL10-CXCR3 axis is involved in the recruitment of peripheral blood human T cells upon CD20-TCB treatment. Despite the strong impact on T cell recruitment, we observed a mild effect of CXCL10 and IFNγ blockade in impairing the efficacy of CD20-TCB in a model of B cell lymphoma in humanized mice. In this regard we could argue that CXCL10 or IFNγ blockade is not sufficient to completely abrogate the efficacy of CD20-TCB mostly because of the contribution of different pathways, such as TNFα induced CXCL9 and CXCL11 [51] in the recruitment of CXCR3^+^ T cells and because of the strong induction of proliferation induced by CD20-TCB. However, T cell recruitment strongly correlates with efficacy demonstrating the importance of recruitment to the TCB MoA. Additional studies are needed in order to identify all the mechanisms involved in human T cell recruitment and how to use them to control tumor growth and potentially develop new immune-therapeutics.

In conclusion, we correlate the quality of the synapse mediated by the T cell bispecific antibody CD20-TCB (RO7082859) with the ability to kill target cells and promote T cell recruitment. We used an innovative DLBCL *in vivo* imaging model using last generation humanized mice that allowed the *in vivo* visualization of the human immune system. We observed that CD20-TCB mediated lymphoma/T cell can trigger the production of IFNγ, which in turn leads to production of CXCL10. This pathway is responsible for increasing T cell retention and recruitment within the tumor microenvironment, supporting the efficacy of the therapy by turning a not-infiltrated tumor into a highly inflamed one. This dataset supports the idea of using such therapeutic classes in the clinics also for tumors which that show poor T cell infiltration.

## Compliance with Ethical Standard

All the experiments were performed according to committed guidelines (GV-Solas; Felasa; TierschG). Experimental study protocol was reviewed and approved by local government (ZH193/2014 and ZH222-17) and by the Ethical Committee of the Roche Innovation Centre of Zurich.

## Acknowledgements

The authors wish to thank all the Roche Innovation Centre Zurich, the colleagues from pREDi for all their support as well as the protein Engineer group especially Claudia Ferrara Koller, Anne Freimoser-Grundschober and Martina Carola Birk.

